# iss-nf: A Nextflow-based end-to-end *in situ* sequencing decoding workflow

**DOI:** 10.1101/2025.10.16.682795

**Authors:** Nima Vakili, Sebastián González-Tirado, Nils Kurzawa, Dmytro Dvornikov, Zeinab Mokhtari, Frank Wippich, Giovanna Bergamini, Rainer Pepperkok, Christian Tischer, Luis A. Vale-Silva

**Author notes:** These authors contributed equally to this work (Nima Vakili, Sebastián González-Tirado, Nils Kurzawa). Heidelberg University, Faculty of Medicine, Heidelberg University Hospital, Institute for Computational Biomedicine, Heidelberg, Germany (Sebastián González-Tirado).

## Abstract

*In situ* sequencing (ISS) offers a powerful approach for spatially resolved gene expression profiling within tissue samples, but the complexity of analyzing the resulting data has limited its broader use. Here, we present iss-nf, a Nextflow-based, end-to-end workflow designed to streamline the decoding of large ISS datasets. The workflow automates critical steps, including image registration, fluorescent spot detection, transcript decoding, and quality control (QC), offering a scalable, reproducible, and user-friendly solution for ISS data analysis. We successfully applied iss-nf on multiple large datasets, including publicly available mouse brain and breast cancer tissue datasets, as well as an in-house non-small cell lung cancer (NSCLC) ISS dataset. The workflow is designed to be accessible for both experienced researchers, as well as newcomers to spatial transcriptomics, providing a robust tool for analyzing large-scale ISS data. Our results suggest that iss-nf is a valuable contribution to the growing field of spatial transcriptomics, enabling precise, modular, reproducible, and, by means of automated tiling and parallelization, scalable analysis of tissue-specific gene expression.

## Introduction

The ability to analyze gene expression in spatial contexts has opened new avenues for understanding the complex interplay of cells within their native environments (Marx, 2021). The rapidly growing field of spatial transcriptomics is propelled by two complementary technological streams: sequencing-based and imaging-based methods (Asp *et al*., 2020; Larsson *et al*., 2021). While sequencing-based approaches such as 10x Visium and Slide-seq (Rodriques *et al*., 2019; Stickels *et al*., 2021) offer transcriptome-wide coverage, they are limited in resolution or sensitivity, and cost-effectiveness. In contrast, imaging-based technologies—including multiplexed fluorescence *in situ* hybridization (FISH) methods like MERFISH (Moffitt *et al*., 2016), seqFISH (Eng *et al*., 2019), and *in situ* sequencing (ISS) (Ke *et al*., 2013)—achieve single-cell or subcellular resolution, often at lower cost, but at the expense of targeting only a subset of the transcriptome.

Among imaging-based spatial methods, ISS, and particularly its hybridization-based variant HybISS (Gyllborg *et al*., 2020), have emerged as powerful tools for targeted spatial transcriptomics. HybISS enables robust detection of nucleic acid sequences in fixed tissue sections using cyclic multicolor fluorescence imaging. Its barcode-based transcript encoding reduces false positives and allows for high levels of multiplexing, as the number of detectable transcripts exponentially scales with the number of imaging rounds. This technique has been successfully used to profile spatial gene expression in diverse biological contexts, including the mouse brain (Qian *et al*., 2020) and human breast cancer (Guo *et al*., 2022). Hybridization-based ISS strategies also underpin commercial platforms such as 10x Genomics’ Xenium, which have been applied to spatially profile gene expression in human and mouse tissues using proprietary chemistries and pipelines (10x Genomics, Xenium *In Situ* Platform: Technical Overview).

Despite its growing adoption, the downstream computational analysis of ISS data remains a significant challenge. Raw ISS data consists of high-dimensional, multi-round, multi-channel image stacks that require careful registration, spot detection, decoding, and filtering. Existing tools such as Starfish (Axelrod *et al*., 2021), the ISS Decoding Tool (Codeluppi *et al*., 2018), and InseqR (Littman *et al*., 2021) provide partial solutions but often lack the scalability, modularity, or full automation required for efficient, reproducible workflows. Many pipelines require substantial manual parameter tuning and are not optimized for large-scale or cloud-based analysis, limiting their applicability to relatively small datasets and making them hard to use for non-expert users.

To address these challenges, we developed “iss-nf”, an end-to-end processing workflow built on Nextflow (Di Tommaso *et al*., 2017) to handle all key steps of ISS analysis: image registration, spot detection with automated thresholding, image tiling, parallelized decoding, and generation of quality control reports. Designed for flexibility, scalability, and reproducibility, iss-nf enables efficient processing of ISS datasets across platforms and scales. We demonstrate the utility of our workflow on public mouse brain and breast cancer datasets, as well as on in-house NSCLC HybISS dataset, highlighting improvements in image registration, decoding accuracy, runtime performance, and usability. Our contribution provides a critical computational resource for researchers adopting *in situ* sequencing in spatial transcriptomics studies.

## Results

### A NSCLC single cell spatial transcriptomics dataset

Lung cancer is the leading cause of cancer-related mortalities with more than two million deaths per year (Torre *et al*., 2015). Despite recent advances in molecular diagnosis and targeted therapies, over half of lung cancer patients die within one year of diagnosis with the 5-year-survival being only 18% (Rahal *et al*. 2025). Lung cancer tumor microenvironment (TME) plays a crucial role in predicting treatment outcomes and patient survival (PMID: 30532012). To profile the TME and underlying gene expression of tumor, stromal and immune cells we designed a target gene panel (n = 242, Supplementary Table 1) with the company Cartana (Stockholm, Sweden), which is part of 10x Genomics since 2020, capturing relevant cell types, phenotypes and genes previously reported to be involved in disease onset, progression and therapy resistance. We imaged n = 7 commercial adenocarcinoma samples using six rounds of imaging.

**Table 1.** Decoded results of seven in-house ISS datasets processed using the ISS-NF pipeline.

| ISS sample code | # Decoded |  | # Detected |  | % Decoded |  | FDR % |  |
| --- | --- | --- | --- | --- | --- | --- | --- | --- |
|  | Postcode | Starfish | Postcode | Starfish | Postcode | Starfish | Postcode | Starfish |
| ISS_1 | 3,197,981 | 2,364,924 | 5,281,263 | 5,281,263 | 60.55 | 44.77 | 0.94 | 6.08 |
| ISS_2 | 5,138,022 | 3,128,118 | 11,488,466 | 11,488,466 | 44.72 | 27.22 | 2.34 | 13.31 |
| ISS_3 | 4,089,345 | 2,426,889 | 6,356,822 | 6,356,822 | 64.33 | 38.17 | 1.29 | 16.93 |
| ISS_4 | 4,111,476 | 2,665,238 | 6,160,360 | 6,160,360 | 66.74 | 43.26 | 1.84 | 8.09 |
| ISS_5 | 3,849,642 | 2,576,200 | 5,624,516 | 5,624,516 | 68.44 | 45.8 | 1.64 | 8.36 |
| ISS_6 | 3,773,360 | 2,359,612 | 5,629,869 | 5,629,869 | 67.02 | 41.91 | 1.37 | 13.94 |
| ISS_7 | 2,106,732 | 515,165 | 4,122,199 | 4,122,199 | 51.1 | 12.49 | 3.4 | 32.01 |

### Challenges with registration of ISS input images using existing workflows

To decode transcripts with their spatial coordinates, we used the Starfish framework (Axelrod *et al*., 2021) with translation-based registration. While we were able to execute the entire workflow, we only retrieved few transcripts per sample and realized that the transcript detection was even worse for samples with significant deformations between imaging rounds (data not shown). This prompted us to develop our own ISS decoding pipeline as a Nextflow workflow based on Starfish, but with support for more advanced registration methods and additional features for workflow automation.

### Nextflow-based workflow

In this paragraph we give a brief overview of our iss-nf workflow; details can be found in the Methods section. To account for the heterogenous outputs of different microscope software and user preferences, we designed iss-nf in a modular way such that it can start from different processing stages (Figure 1). The input requirements for running iss-nf are anchor nuclei and spots as well as nuclei and spots images from all imaging rounds. The nuclei images of the rounds are used to determine the spatial deformation of each round by registering each of them to the “reference” anchor nuclei image, which is acquired at the beginning of the experiment. The anchor spots image contains the coordinates of all transcripts, while the rounds spots images are multiple fluorescence channels and encode the transcript genes. In addition, the workflow requires a codebook with the barcodes for individual transcripts and config files with adapted parameters to the respective experiment, e.g. number of imaging rounds, file paths, and names of targets with empty barcodes. Our workflow starts with stitched input images in TIFF format which are then passed to a registration module that applies translational, Euler, affine and b-spline transformations to achieve single pixel-accuracy registration of images from rounds (Ntatsis *et al*., 2023). Next, registered images are automatically tiled such that the subsequent spot detection can be run in parallel and thereby speed up the analysis. Decoding is performed with Starfish (Axelrod *et al*., 2021) or PoSTcode (Gataric *et al*., 2021). Last, a quality control report is generated which also includes the computation of a false discovery-rate (FDR) per sample, utilizing the user provided empty barcodes (false negatives).

**Figure 1.**
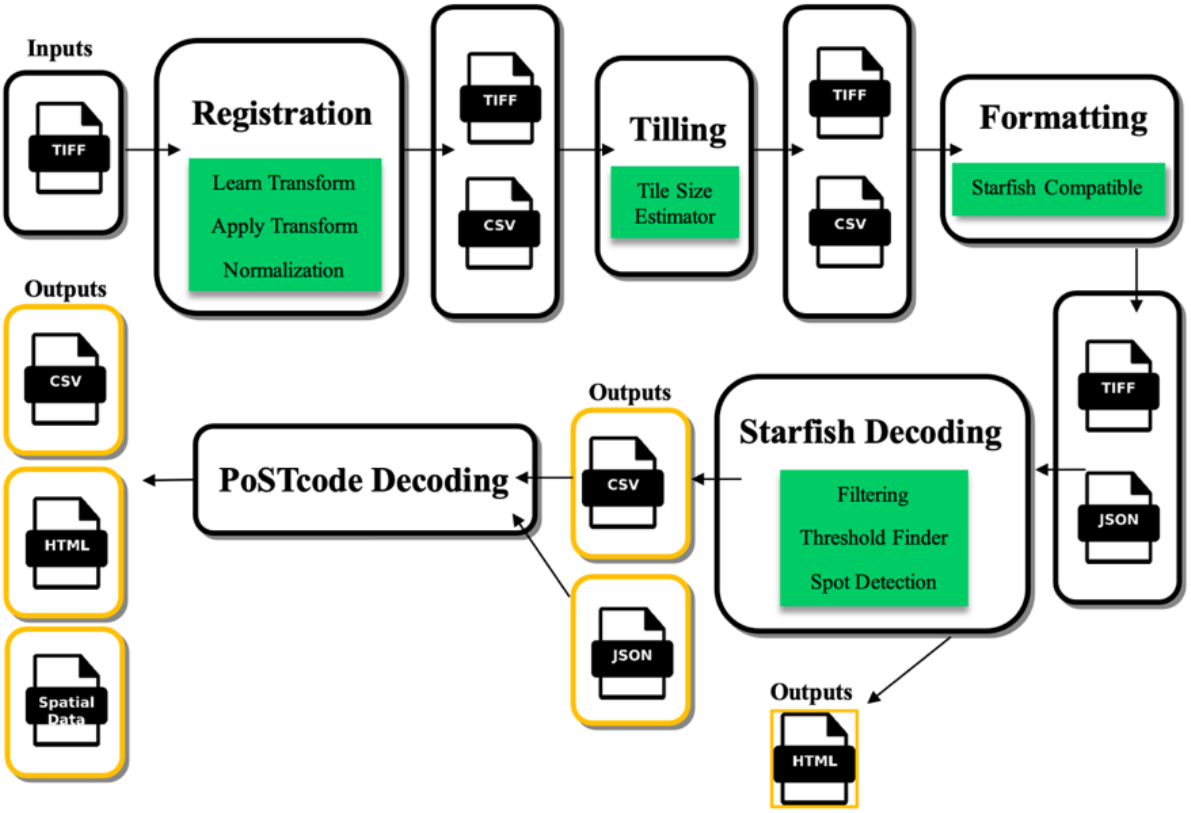
Workflow diagram of the ISS-NF pipeline. The process begins with the Registration module, which expects input TIFF images. Next, the data undergoes Tiling for improved computational and memory efficiency, followed by a Format conversion to make it compatible with Starfish. Next, a decoding approach using the PoSTCode decoding method is applied. Decoding using the PerRoundMaxChannel Starfish method is also supported (but not recommended). The final output consists of a CSV table containing all decoded transcripts, along with an HTML quality control report summarizing metrics and images from all modules. Additionally, a SpatialData output is generated, allowing users to conveniently visualize results in Napari.

Using iss-nf to process the same in-house NSCLC samples yielded much improved results (Figure 2). To further evaluate iss-nf, in addition to applying it to the in-house NSCLC samples, we also validated it by analyzing two publicly available ISS datasets, comparing our results to the published decoding results.

**Figure 2.**
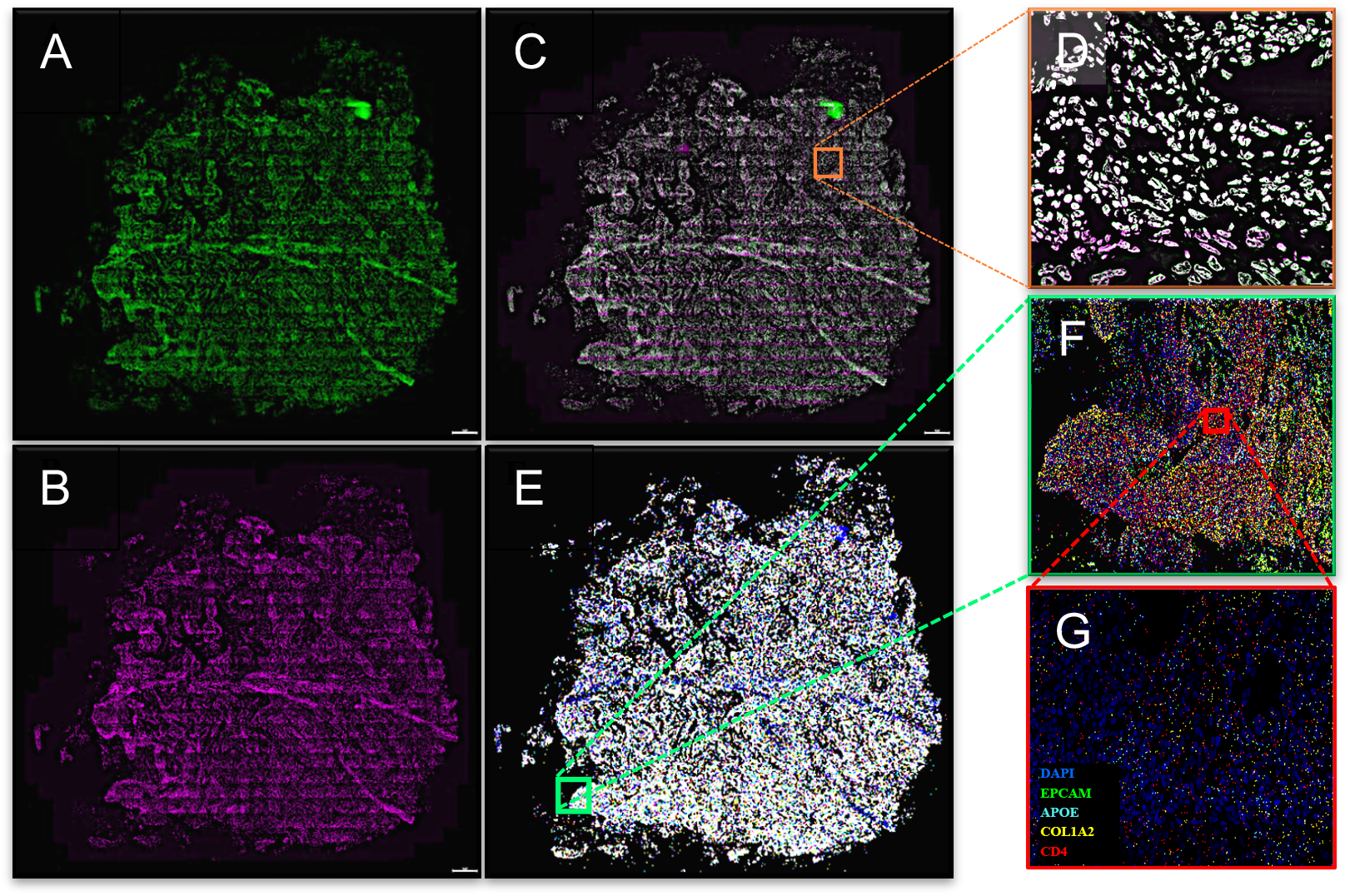
Results of the ISS-NF pipeline applied to in-house in situ sequencing data. (A) Anchor DAPI staining (green). (B) DAPI staining from Round 1 (magenta). (C) Overlay of (A) and (B), demonstrating the accuracy of the registration process (white areas indicate perfect matching of green and magenta signals). (D) Zoom-in of a region of interest (ROI) from (C), highlighting precise registration. (E) Decoded transcripts overlayed on DAPI-stained nuclei (blue). (F) Zoom-in of a selected ROI from (E), highlighting the expression of four selected genes. (G) Further zoom-in of the ROI in (F) for detailed visualization of gene expression patterns.

**Figure 2.**
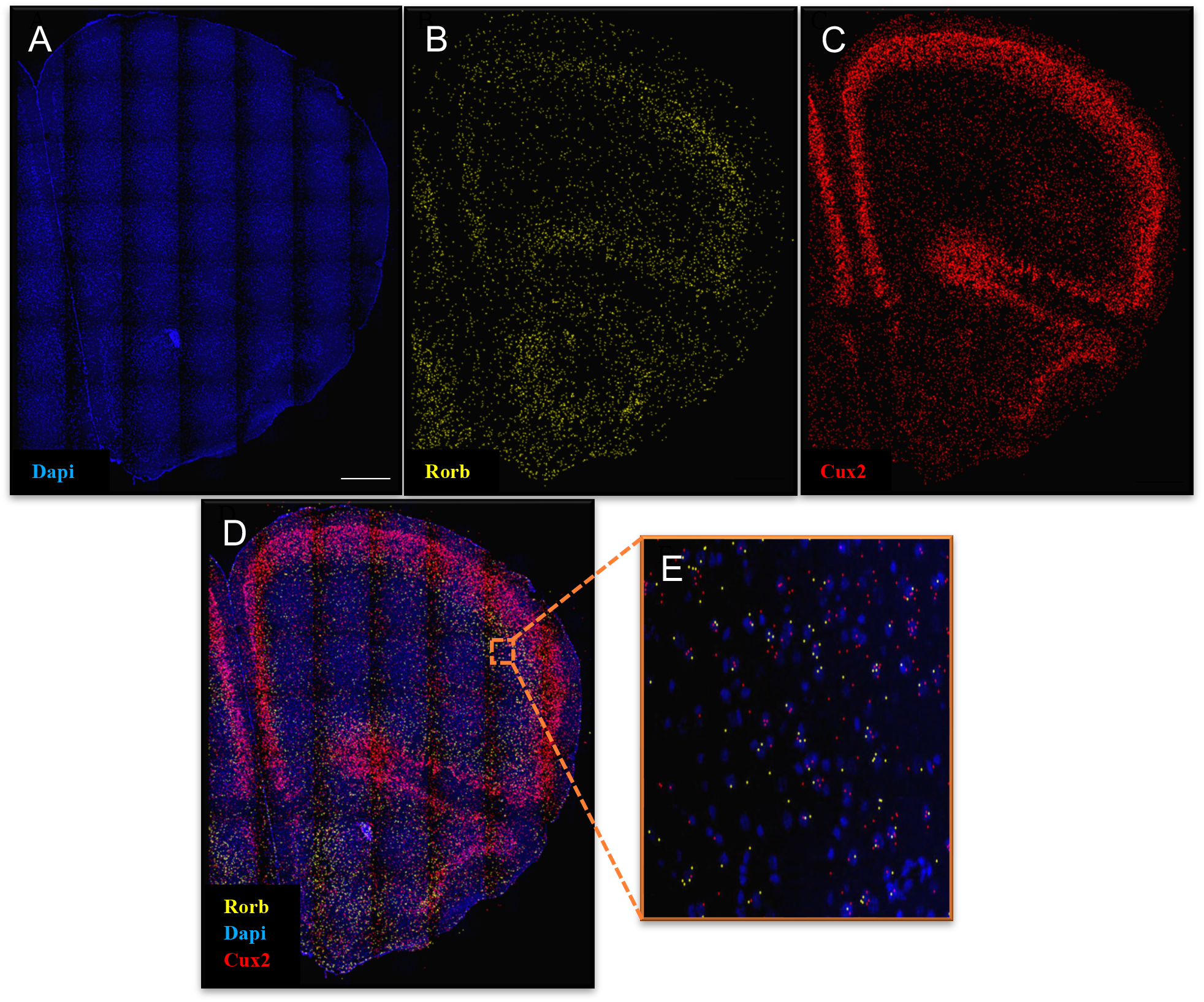
iss-nf results for the mouse brain ISS dataset by Gataric et al. (A) DAPI staining in blue. (B) Decoded spatial pattern of the Cux2 gene (GTCC barcode) in green. (C) Decoded spatial pattern of the Rorb gene (GCGG barcode) in red. (D) Overlay of (B) and (C) on DAPI-stained nuclei (blue), showing the spatial distribution of Cux2 and Rorb. (E) Zoomed-in section of (D) for a detailed view of the spatial patterns of the two genes.

### Evaluation of the iss-nf workflow on a mouse brain dataset

First, we chose to analyze the mouse brain ISS dataset (Gataric, Bayraktar, & Gerstung, 2021a) published along with the PoSTcode method. The mouse brain has a clear structure and is well studied, e.g. by methods such as *In Situ* Hybridization (ISH), such that spatial expression patterns can be easily contextualized. This aids the comparison of our results with published results. The size of this dataset is 11 GB and, utilizing high-performance computing (HPC; resources managed automatically by Nextflow), the analysis was completed within 12 minutes. Our iss-nf workflow achieved an overall number of 384,737 detected spots, a decoding percentage of 81.9 and an FDR of 0.16%. These results mark a minor 1.3% improvement in terms of decoded spots compared to the results reported by Gataric (Gataric *et al*., 2021b). The spatial expression patterns of the genes *Ror* and *Cux* (Figure 3) were also comparable to the published ISS and ISH results. This demonstrates that our workflow recapitulates previously validated results.

### Evaluation of the iss-nf workflow on a human breast cancer dataset

Next, we chose to process the breast cancer dataset by Lomakin (Lomakin *et al*., 2022) to test our workflow on a large human dataset. The size of this dataset is 88 GB and the analysis was completed within 2 hours. We achieved an overall FDR percentage of 0.68% and comparable or higher numbers of decoded spots for most true probes and lower numbers of decoded negative control probes than in the original publication (Figure 4). Spatial patterns of individual probes are harder to contextualize in tumor samples compared to the mouse brain example; however, the decreased fraction of decoded negative control probes lead us to conclude that our method is obtaining specific results and the increase in decoding of several probes indicates high sensitivity.

**Figure 4.**
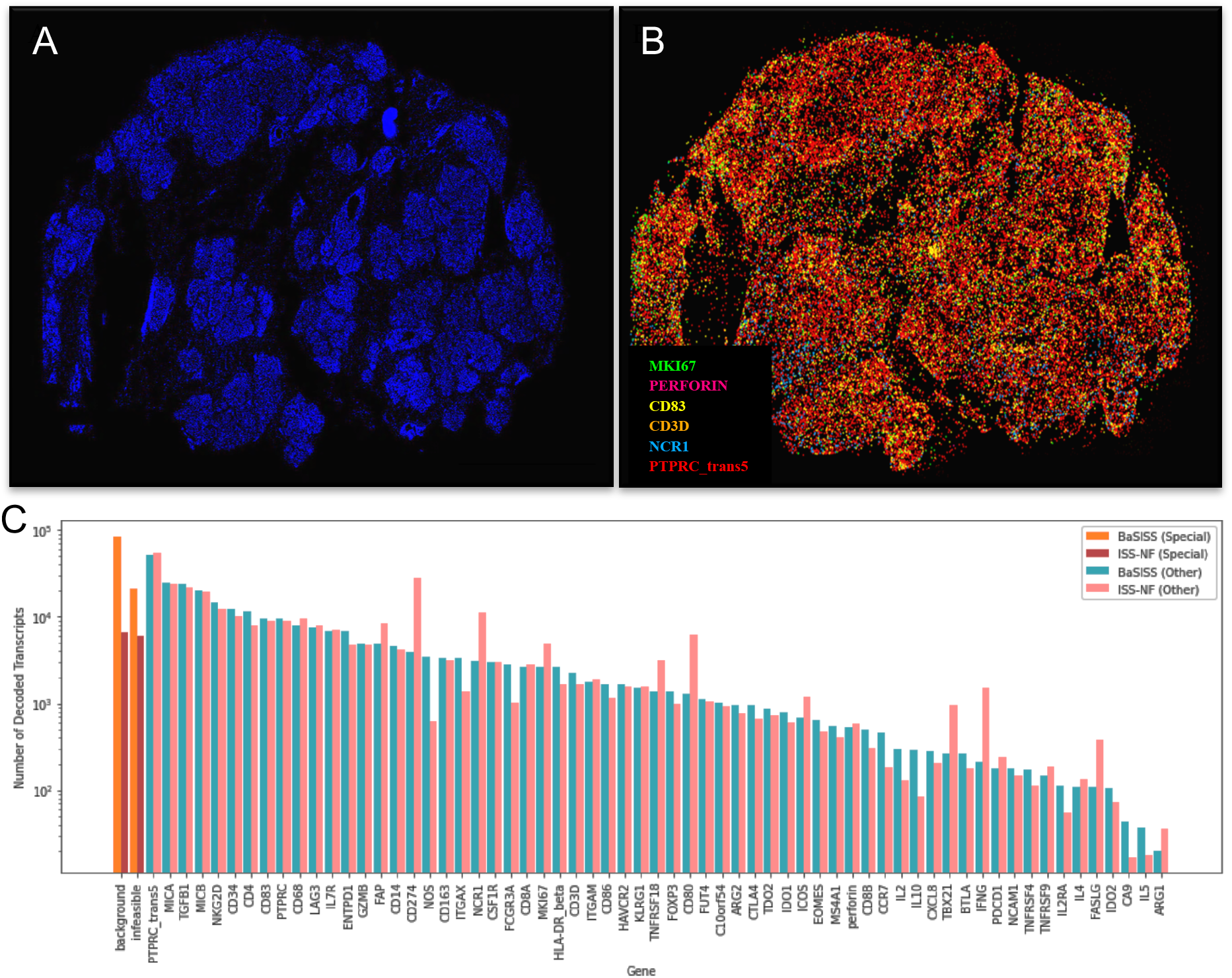
Pipeline results of a human breast cancer ISS dataset by Lomakin et al. Comparison of decoding results between Lomakin et al. and iss-nf. (A) DAPI staining (blue). (B) Decoded transcript maps for six selected genes. (C) Comparison of decoded genes, background, and infeasible signals decoded by the BaSISS method (Lomakin et al., 2022) and the iss-nf pipeline, illustrating differences in gene decoding accuracy and signal interpretation.

### Analysis of an in-house NSCLC dataset

In addition to validating our workflow on public datasets, we also applied iss-nf to our in-house generated NSCLC HybISS dataset. The size of a typical dataset was 70 GB and the analysis was completed within 5 hours. We achieved high numbers of detected spots (multiple millions per samples; approximately 3-fold more than with the standard Starfish workflow in the case of ISS_4) and FDR percentages below 5% for all samples (Table 1) using PoSTcode for decoding. Next, we further processed the decoded NSCLC samples ISS_1 and ISS_4 by assigning transcripts to cells and annotating cell types. This was done by applying the optimal transport-based TACCO algorithm for annotation of single molecule data (Mages *et al*., 2023) using a single-cell RNA-seq (scRNA-seq) reference dataset. We used the dataset by (Zhang *et al*., 2019) annotated cell types using cell assign and a list of marker genes and then first annotated single molecules and subsequently segmented single cells using TACCO. Obtained cells captured on average 32 transcript counts of on average 18 unique genes (Figure 5A and B). While ISS_4 represented an “excluded” phenotype, characterized by tumor cells isolated from immune cells, ISS_1 showed extensive infiltration of macrophages in tumor regions (Figure 5C and D).

**Figure 5.**
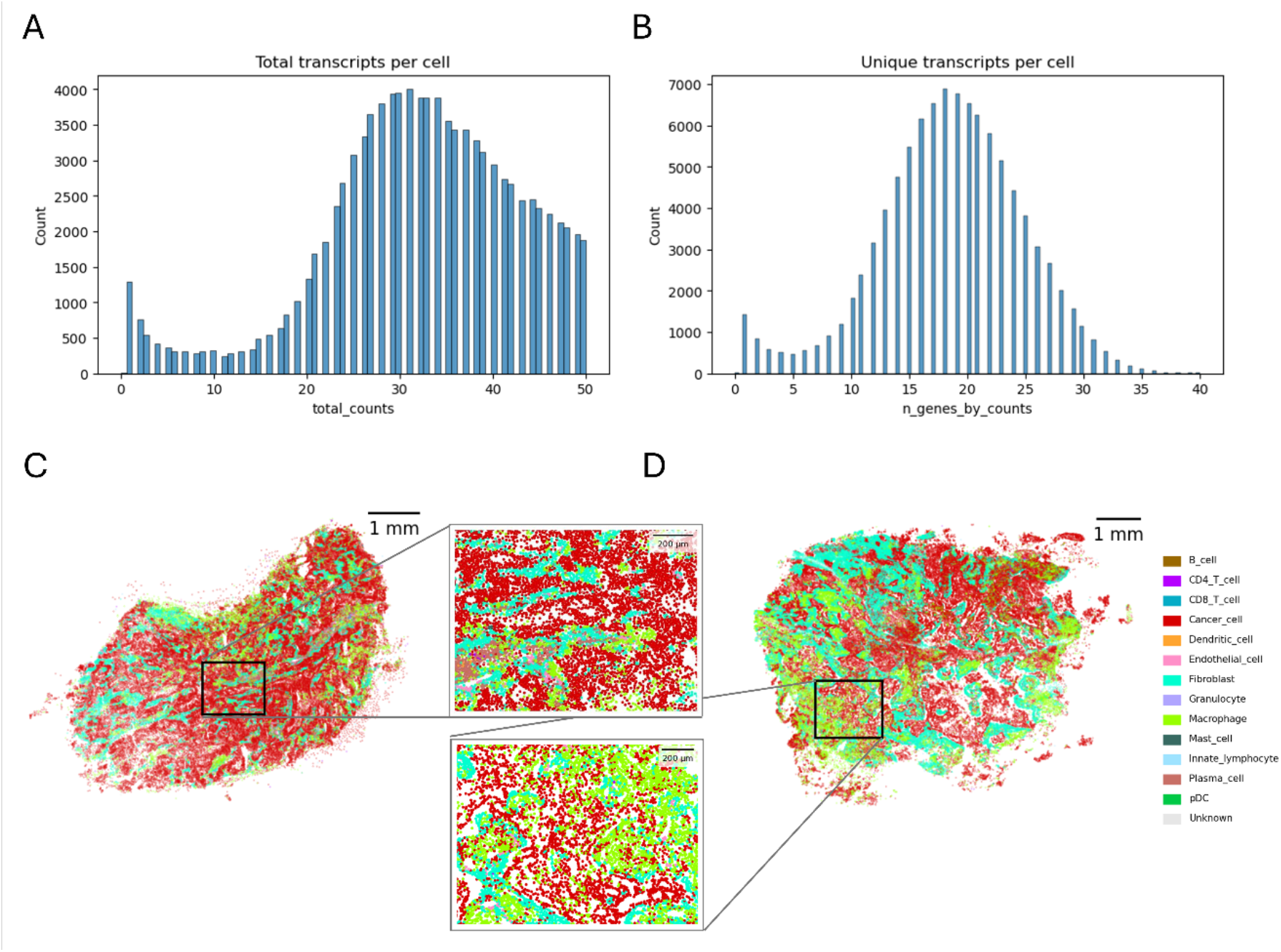
Transcript coverage and cell type maps obtained from exemplary NSCLC samples. Histograms of A) total transcript counts and B) number of unique genes per cell for ISS_4. Cell type maps of C) ISS_4 and D) ISS_1.

## Methods

### Image acquisition

Cryosections (8-um thick) were prepared using Leica cryostat and stored at -20°C until use. All sections were processed for *in situ* hybridization within two weeks of preparation using the Cartana protocol, following the manufacturer’s instructions. During all incubation steps, 100 uL SecureSeal hybridization chambers (GraceBio) were applied to ensure fluid containment and uniform reagent distribution. Imaging was conducted using an Olympus VS200 slide scanner across DAPI, FITC, Cy3, Cy5 and Cy7 channels. Exposure times were optimized per channel and cycle to prevent signal saturation. Mild on-the-fly image deblurring was applied to all non-DAPI channels. A five-plane z-stack was acquired per section and merged into a single projection using Olympus Extended Focal Imaging (EFI) software. Final image datasets were exported as uncompressed BigTIFF files for downstream processing.

### Stitching and preparation of input images

The iss-nf pipeline requires input images to be stitched prior to further analysis. Users can achieve this by utilizing their microscope’s stitching software or external stitching tools. We recommend using Ashlar (Ramos *et al*., 2021), a popular open-source tool for stitching images from cyclic imaging experiments. The input format of all images must be TIFF. A key step in preparation to start the iss-nf pipeline is ensuring a consistent image naming scheme. Images must be named using the format “r{ROUND_INDEX}_{CHANNEL_NAME}.tiff”, where “r” is followed by “ROUND_INDEX”, an integer specifying the imaging round, and “CHANNEL_NAME”, identifying the fluorescence channel. For example, an image from round 1 and channel Cy3 must be named “r1_Cy3.tiff”. Adhering to this naming scheme is essential for the pipeline to accurately associate the correct rounds and channels during the subsequent steps of image registration and decoding.

Image registrationAccurate registration of images from different imaging rounds is crucial for ISS data processing as barcode decoding requires pixel-level precision to correctly associate signals from each imaging round and channel. In iss-nf this is achieved using ITK-Elastix for registration, which employs a multi-step approach: starting with translation, followed by rotation, affine transformation, and finally, b-spline deformation correction. The b-spline component is particularly useful for samples that exhibit distortions during image acquisition. Our registration module applies these transformations to all stitched images, typically achieving pixel-wise alignment across all imaging rounds.

### Image normalization

To ensure consistent thresholding and spot detection for all rounds and channels, each image was percentile (0, 0.98) normalized to values between 0 and 1.

### Automated tile size estimation and tiling

To handle large images and avoid memory issues, automated tiling was implemented. This includes dividing the images into manageable tiles, enabling efficient parallel processing with Nextflow’s parallel processing capabilities. This strategy ensures that memory limitations do not hinder the workflow while optimizing processing time by leveraging distributed computing.

### Automated threshold detection

To ensure consistent thresholding and spot detection for all rounds and channels, stitched and registered images were percentile (0, 0.98) normalized to values between 0 and 1. Laplacian of Gaussian image enhancement was applied prior to spot detection. To automate spot detection, which typically requires manual threshold determination, an automated threshold selection algorithm that minimizes the false discovery rate (FDR; 10x Genomics, Specificity, sensitivity, throughput, and data analysis for *in situ* molecular spatial technologies; details below) while maximizing the number of decoded spots was implemented. The algorithm utilizes the required list of empty barcodes (10-20% of their gene panel size) with a Hamming distance of minimum two to the most similar true barcode, which are used for assessment of false positive decoding. The algorithm selects the three tiles with the highest intensities (after optional top-hat filtering; see below) and then calculates the number of decoded spots and FDR across 10 thresholds ranging from 0.001 to 0.009. For each of the three tiles, the threshold that achieves the best trade-off between decoded spots and FDR is determined; the resulting three threshold values are then averaged to yield the final threshold that is used for the whole image.

### Optional tophat filtering

To further enhance image quality, an optional top-hat filtering step was implemented, which can be enabled via the Nextflow configuration file. This filter reduces uneven background, thereby improving spot detection by increasing the contrast between spots and surrounding tissue. Based on our empirical observations, a disk-shaped structuring element with a radius of 5 pixels provided an effective balance between background reduction and the preservation of local intensity features. This setting was applied consistently across all datasets analyzed in this work.

### Transcript decoding

Following spot detection, transcript decoding is performed using the Starfish package. The pipeline was implemented to support the PerRoundMaxChannel decoding method, which identifies transcripts based on the most intense channel in each imaging round. In addition to decoding with the Starfish framework, the iss-nf pipeline also supports an alternative decoding strategy using the PoSTcode (Probabilistic Image-based Spatial Transcriptomics Decoder) algorithm (Gataric *et al*., 2021). We found PoSTcode results to be consistently superior to Starfish’s PerRoundMaxChannel, particularly when there is variability in signal quality across imaging rounds.

### Quality control and false discovery rate

The iss-nf workflow was designed to incorporate quality control (QC) measures at key stages to ensure high accuracy throughout the process. An intermediate QC step was implemented by providing a QC report in HTML format that quantifies registration quality. This was done to allow evaluation of alignment accuracy and to determine whether to proceed with subsequent modules or adjust registration parameters for improved results. The workflow also outputs a comprehensive general QC report. This includes an HTML file summarizing various statistics, plots, and QC metrics for each Nextflow module. Additionally, a CSV file is produced, detailing all detected and decoded spots, their coordinates, gene annotations, and the percentage of spots passing thresholds for both decoding approaches (Starfish PerRoundMaxChannel and PoSTcode).

To assess the false discovery rate (FDR), a method first proposed by 10x Genomics was implemented (10x Genomics, Specificity, sensitivity, throughput, and data analysis for *in situ* molecular spatial technologies). Briefly, “empty” barcodes, i.e. barcodes designed in the same way as true barcodes but with no corresponding molecular probes in the experimental procedure, are used as negative controls. Total decoding results, including true and “empty” barcodes, are then used to compute the FDR% as follows:

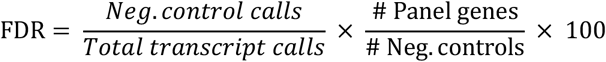

The workflow also outputs data in SpatialData format (Marconato *et al*., 2025), allowing users to inspect and further analyze the data in the python ecosystem, including overlaying all or selected target gene expressions on nuclei images, providing an interactive platform for reviewing results. These outputs collectively offer a thorough assessment of the workflow’s performance and support further downstream analyses.

### Mapping transcript molecules to cells

A NSCLC single-cell RNAseq (scRNAseq) dataset was downloaded, pre-processed according to scanpy default recommendations and annotated for cell types using a marker gene table and the software cellassign (Zhang *et al*., 2019). The optimal transport-based software TACCO with default parameters was then used to map decoded ISS transcript molecules to cells via its segmentation-free method using the scRNA-seq dataset as a reference.

## Conclusion

We present iss-nf, a Nextflow-based pipeline designed for the analysis of *in situ* sequencing (ISS) data. This workflow provides an efficient, scalable, and reproducible solution for spatial transcriptomics research, facilitating the processing of large datasets with minimal computational overhead.

By leveraging containerization and workflow automation, iss-nf ensures consistency across different computing environments while allowing for seamless integration with existing bioinformatics tools. Additionally, its modular design enables customization and adaptability to different experimental setups.

We applied it to two different public ISS datasets and obtained on par or improved results compared to original publication processing results. Successful extraction of biologically meaningful insights from an in-house generated NSCLC ISS dataset further underscores the value of our workflow.

Overall, iss-nf enhances the accessibility and reproducibility of ISS data analysis, empowering researchers to extract deeper insights into spatial gene expression patterns and tissue organization.

## Acknowledgements

We thank Lena Eismann, Kathrin Jansen and Felix Frauhammer for pre-processing of the scRNAseq dataset by Wu *et al*.

## Code availability

The iss-nf pipeline is an open-source workflow designed for scalable and reproducible decoding of *in situ* sequencing data. It is freely available on GitHub at https://github.com/BioImageTools/iss-nf under an open-source license. To help users get started, we provide a publicly available half-mouse-brain ISS dataset on Zenodo (https://zenodo.org/records/14884160). This dataset can be used to test and evaluate the pipeline’s functionality. Comprehensive documentation, example datasets, and step-by-step tutorials guide users through installation, configuration, and execution. The workflow is modular and adaptable to different experimental designs, and community contributions are encouraged to extend its capabilities.

## Data availability

Mouse brain raw data is available on the BioImage Archive (https://www.ebi.ac.uk/biostudies/bioimages/studies/S-BSST700). In addition, we provide the same data formatted to work with our iss-nf workflow on Zenodo (https://zenodo.org/records/14884160). Breast cancer raw data is available on the BioImage Archive (https://www.ebi.ac.uk/biostudies/BioImages/studies/S-BIAD537?query=S-BIAD537). Seven in-house NSCLC sample datasets have been made publicly available via the BioImage Archive (https://www.ebi.ac.uk/biostudies/bioimages/studies/S-BIAD2277).

## Computational Environment and Workflow Execution

All computational analyses were conducted on a HPC cluster using the Slurm job scheduler. The iss-nf pipeline was orchestrated using Nextflow, enabling scalable and reproducible execution of modular decoding tasks. Custom Nextflow configurations were defined for each process using task-specific labels, which specify computational resource requirements (e.g., CPUs, memory, modules) and Slurm directives. Data staging and loading were optimized for efficient parallel execution. All configuration profiles used for Slurm execution are available in the pipeline repository for reproducibility.

